# CovidExpress: an interactive portal for intuitive investigation on SARS-CoV-2 related transcriptomes

**DOI:** 10.1101/2021.05.14.444026

**Authors:** Mohamed Nadhir Djekidel, Wojciech Rosikiewicz, Jamy C. Peng, Thirumala-Devi Kanneganti, Yawei Hui, Hongjian Jin, Dale Hedges, Patrick Schreiner, Yiping Fan, Gang Wu, Beisi Xu

## Abstract

Infection with severe acute respiratory syndrome coronavirus 2 (SARS-CoV-2) in humans could cause coronavirus disease 2019 (COVID-19). Since its first discovery in Dec 2019, SARS-CoV-2 has become a global pandemic and caused 3.3 million direct/indirect deaths (2021 May). Amongst the scientific community’s response to COVID-19, data sharing has emerged as an essential aspect of the combat against SARS-CoV-2. Despite the ever-growing studies about SARS-CoV-2 and COVID-19, to date, only a few databases were curated to enable access to gene expression data. Furthermore, these databases curated only a small set of data and do not provide easy access for investigators without computational skills to perform analyses. To fill this gap and advance open-access to the growing gene expression data on this deadly virus, we collected about 1,500 human bulk RNA-seq datasets from publicly available resources, developed a database and visualization tool, named CovidExpress (https://stjudecab.github.io/covidexpress). This open access database will allow research investigators to examine the gene expression in various tissues, cell lines, and their response to SARS-CoV-2 under different experimental conditions, accelerating the understanding of the etiology of this disease to inform the drug and vaccine development. Our integrative analysis of this big dataset highlights a set of commonly regulated genes in SARS-CoV-2 infected lung and Rhinovirus infected nasal tissues, including OASL that were under-studied in COVID-19 related reports. Our results also suggested a potential FURIN positive feedback loop that might explain the evolutional advantage of SARS-CoV-2.

## INTRODUCTION

Infection with severe acute respiratory syndrome coronavirus 2 (SARS-CoV-2) in humans could cause Coronavirus disease 2019 (COVID-19). Since it was first discovered in Dec 2019, SARS-CoV-2 has become a global pandemic that spread into 192 countries, infected approximately 163 million people and caused 3.3 million of deaths to date (2021 May) (Dong *et al*). The scientific community quickly responded to COVID-19 and public data sharing in health crises (Littler *et al*) played a critical role across all aspects of combatting against SARS-CoV-2. For instance, Nextstrain and GISAID databases allow sharing and comprehensive real-time analysis to monitor the virus evolution and adaptation (Hadfield *et al*; Shu & McCauley) since the first sequencing data were available (Wu *et al*). MOBS LAB and MIDAS enabled the epidemic spread modeling that facilitated policy makers to draft emergent regulations (Chinazzi *et al*; Kenah; Kucharski *et al*; Sun *et al*). Platforms like Vivli shed a light on sharing clinical trial data anonymization to accelerate scientific progress (Li *et al*). The structure sharing on Protein Data Bank (Burley *et al*; Jin *et al*; Walls *et al*) and CoV3D (Gowthaman *et al*) have been instrumental for investigators to understand SARS-CoV-2’s function and mechanism. Databases that document drug repurposes were developed either based on 3D structure (Chen *et al*) or literature searching (Tworowski *et al*). Web portals such as LitCovid (Chen *et al*, 2020, 2021a) and COVIDScholar (Trewartha *et al*) were developed to reduce the time for acquiring information, while Cochrane were more focused on collecting clinical trial studies (Hilton *et al*). The gene sets collected by PAGER-CoV (Yue *et al*) and COVID-19 Drug and Gene Set Library (Kuleshov *et al*) could be used for cross-examining different results and prioritization of potent drugs.

On the other hand, the fast growth of COVID-19 related publications (Palayew *et al*) led to many cases of unreliable results, some of which were unfortunately retracted (Abritis *et al*; Else, 2020). Thus, re-analysis and critical curation of the ever-growing available data in an unbiased manner becomes a necessity to ensure the reproducibility of scientific conclusions and further propel advances toward therapeutics. Of the many published studies depositing raw data in public repositories including GEO (Barrett *et al*), bulk RNA-seq datasets have taken the largest share. However, few databases enable rapid and easy access to such as quantitative details (Cantelli *et al*; Ziegler *et al*) in RNA-seq, and none provides analysis functions for world-wide investigators without programming skills.

Thus, we sought to develop an open-access database so investigators could easily examine the comprehensive, consistently re-processed data by performing basic analysis with intuitive, customized, and instant visualization. Our main goals are to facilitate open-access to publicly available COVID19 related bulk RNA-sequencing data, to provide standardized results for investigators to use to generate hypotheses for subsequent genetic, biochemical, and functional experimental analyses, and to present rich visualization tools so investigators can make publication-ready figures to accelerate publication process. With this effort, we have curated and analyzed up to date the largest set of 1,468 SARS-CoV-2 related bulk RNA-seq samples from 43 independent studies. Those data, after quality control, were also used to run a total of 315 contrasts of differentially expressed genes analyses, and utilizing several alternative postprocessing approaches, ultimately allowed for identification of 20,036 differential SARS-CoV-2 related gene signatures.

## RESULTS

### Overview of Data Processing Workflow for CovidExpress

**Figure 1A** describes the general schema of the data curation and analysis. We collected human RNA-seq data by using the query “((covid-19 OR SARS-CoV-2) AND gse[entry type]) AND “Homo sapiens”[porgn:__txid9606]” at NCBI Gene Expression Omnibus(GEO) website. From 43 studies (9 published, **Supplementary Table S1**), we collected 1,468 samples (4.7TB raw data) from total representing 227 experimental groups. We integrated quality control (QC) code from RSeQC (Wang *et al*) so that optimal parameters for processing pipeline were automatically chosen (see Material and Methods) based on the strand information of RNA-seq protocol (**Supplementary Figure S1A, S1B**). Next, we reviewed the quality control metrics and found that most data had good sequencing depth (**Supplementary Figure S1C**) and mapping rate (**Supplementary Figure S1D**). Nevertheless, the genomic distribution of the sequencing reads revealed that about 21.2% of the samples were biased toward intronic reads (**Supplementary Figure S1E, Supplementary Table S1**, Material and Methods), we reasoned that these biases could be explained by protocol differences and experimental variations. We still included those samples since our processing method was only counting exonic reads as for gene level expression quantification. Also, these biases did not affect our differential expression analysis since we would only compare samples within the same study cohort.

**Figure 1.**
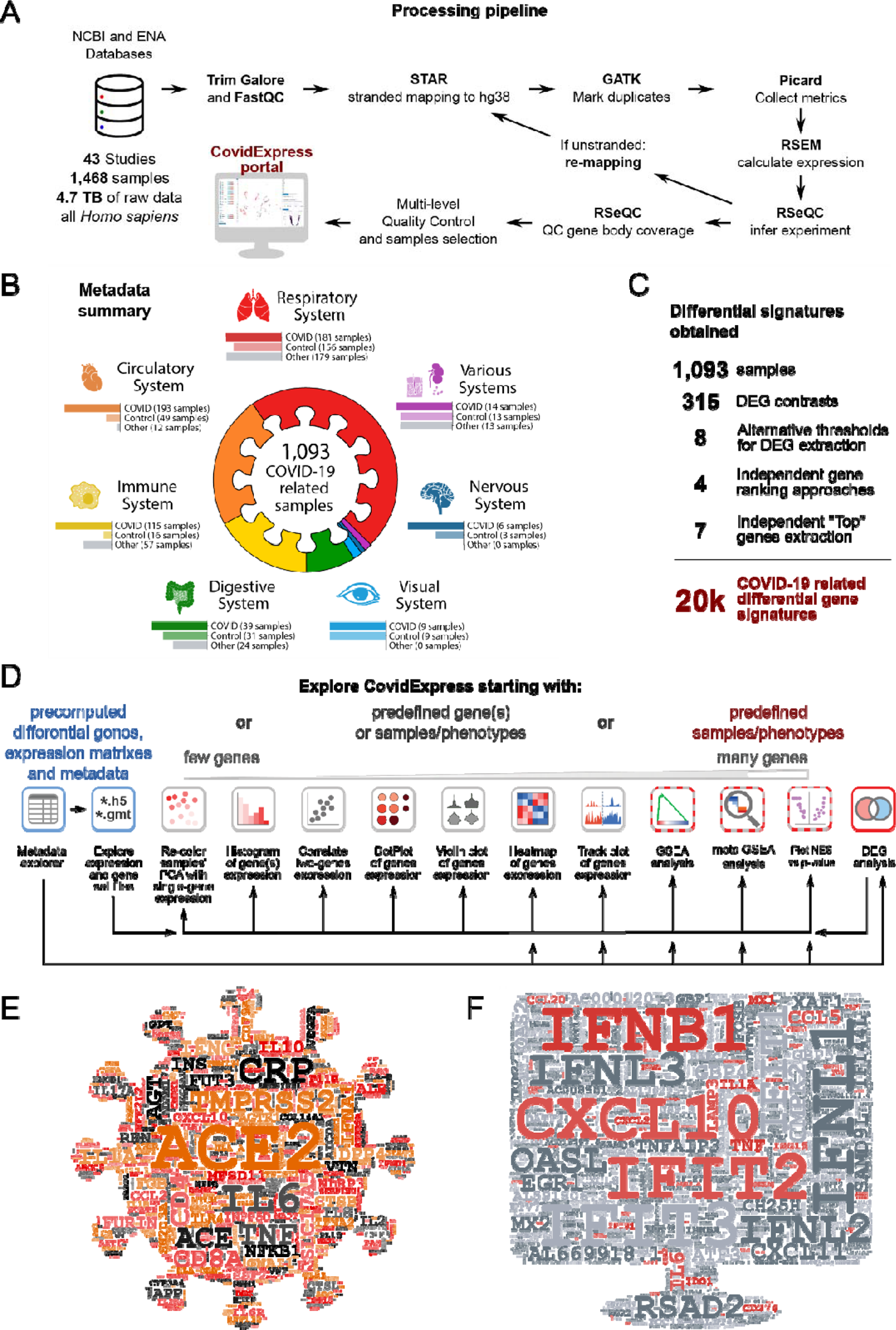
Overview of RNA-seq processing Workflow. **A** General schema of data collection and processing. See Material and Methods part for details. **B** Overview of samples distribution for finalized samples used. **C** Summary of the number of samples, DEG contrasts, differential gene signatures extraction approaches and final number of differential gene signatures. **D** Overview of general approaches for CovidExpress exploration, and analytical and visualization tools that may be utilized within the portal. **E** Word cloud highlighted genes frequently reported in literatures related to SARS-CoV-2. **F** Word cloud highlighted genes frequently shown in top 500 regulated genes from this study(315 contrasts), genes also in Figure 1E were colored in red.

We noticed that several studies were strongly enriched with sequences from 3 UTRs that would affect our analyses. We found that this 3 UTR bias could be explained by experimental protocols. For instance, a study from GSE154936 employed QuantSeq (Moll *et al*) protocols known by its 3’UTR bias in order to utilize the low input or low quality RNA materials. To systematically identify similar samples, we aggregated the normalized gene-wise binned coverage for each sample and used hierarchical clustering to separate datasets into 9 clusters by gene-coverage similarity (**Supplementary Figure S1F)**. Principal component analysis (PCA) (Lever *et al*) of the gene-body coverage profiles showed that clusters 1, 2 and 3 were distinct from the other clusters (**Supplementary Figure S1G)**. A summarized plot confirmed that clusters 1, 2 and 3 were strongly biased toward the 3 UTR (**Supplementary Figure S1H**).

Clusters 4, 5 and 6 also shown a slight 3’UTR bias, but we decided to keep them since they achieved good coverage for gene body (**Supplementary Figure S1H**). Together, we included data from clusters 4-9 as the finalized sample list for our database and all downstream analyses. All our QC metrics are listed in **Supplementary Table S1**. Our finalized sample list includes 1093 samples, where 557 (50.96%) were of samples infected with SARS-CoV-2. Most samples were from Respiratory, Circulatory, and Immune systems (**Figure 1B, 1C**). An overview of the overlap between different analytical criteria were summarized in **Supplementary Figure S1I and S1J**.

To help investigators begin their own analyses using CovidExpress portal (**Figure 1D**), we took two approaches to highlight the currently known important genes in COVID-19 research. We first used machine learning-based tool for retrieving bioconcept annotations from literature research. We extracted gene annotations from 108,627 COVID-19 related literature sources, and weighed genes by their appearance frequency in Pubtator annotation (Wei *et al*, 2019) (**Figure 1E**). Per our expectation, *ACE2* and *TMPRSS2* emerged as the top studied genes (Clausen *et al*; Hoffmann *et al*, 2020; Liu *et al*, 2020; Ziegler *et al*., 2020). Many other top genes were also very interesting, such as C-reactive protein (CRP), which may be an early marker of COVID-19 (Luo *et al*, 2020). Several other top genes require further genetic, biochemical, and functional characterization using follow-up experiments.

Next, we extracted metadata using GEOmetadb and manually reviewed data for accuracy (Zhu *et al*). We performed a total of 315 differential expression (DE) genes analyses between groups for each study, and we asked which genes frequently showed up as top differentially expressed shared among all our comparisons (**Figure 1C and 1F**). We found that many COVID-19 context studies reported that these top differentially expressed genes were important (**Supplementary Table S2**). Some of the top genes that frequently appeared in literature research approach such as *IL6, IL1A* and *TNF*, were also found among the top differentially expressed genes. Interestingly, top genes in literature-based search, such as *ACE2*, *TMPRSS2*, and *CRP*, were not among the top differentially expressed genes. This suggests that although it have been repeatedly reported that SARS-CoV-2 need ACE2/TMPRSS2 to infect cells (Clausen *et al*.; Hoffmann *et al*., 2020; Liu *et al*., 2020; Ziegler *et al*., 2020), *ACE2/TMPRSS2* expression level was not elevated following infection. Thus, the most dramatic differential expression observed in RNA-seq was more related to the innate immune defense mechanism. In support of this notion, many interferon genes and inflammatory cytokine and chemokine genes were frequently found as top differentially expressed genes (**Figure 1F**). The results of such meta-analysis itself has a power to guide future molecular studies to determine the functional impact of these genes and the resulting proteins in the disease pathogenesis. Overall, our RNA-seq analyses pointed to the innate immune defense mechanism as the most differentially regulated following SARS-CoV-2 infection.

### CovidExpress Web Portal Overview and Key Functional Components

Large datasets could be challenging to explore especially for investigators without programming skills. Thus, we built our server blueprinted from cellxgene interface, which is a tool that was originally designed for exploring single-cell RNA-seq data and comes with a rich set of features (Cakir *et al*). Furthermore, we added designed features and extended the functional set of cellxgene components to allow investigators to visualize and customize their results as well as to run more in-depth analyses (**Figure 2A**). Beside all the functionalities cellxgene supports, our server further allows investigators to visualize the expression of their genes of interest and exports publication-ready figures such as violin plots, dot plots, track plots and heatmaps (**Figure 1D, 2B, Supplementary Figure S2A, S2B).** Importantly, we developed additional analytical components so that investigators could explore their own gene sets (i.e. lists of genes of interest). Our server enables investigators to calculate enrichment scores against all our pre-computed regulated gene ranks by using Gene Set Enrichment Analysis (GSEA) (Fang; Subramanian *et al*). The results will be presented to investigators in an interactive manner so they can select the top comparisons of interest (**Figure 2C**). To ensure the reliability of the results, the codes for calculation scores for GSEA have been carefully reviewed. We implemented a procedure for multiple hypothesis testing: p-values were controlled by False Discovery Rate (FDR) from all pre-computed p-values for each comparison. This would allow investigators to assess the specificity of their gene set comparison to gene sets from other databases including MSigDB (Liberzon *et al*) and COVID-19 related gene set databases we or the investigators curated (Kuleshov *et al*.). Finally, the investigators can review the strength and confidence of their gene set enrichment by examining the output volcano (**Figure 2D**) and GSEA plots (**Figure 2E**). The investigators could also choose to explore our pre-computed GSEA results from different gene set databases (**Figure 2F**), using different gene-ranking strategies (**Supplementary Figure S2C**), or based on different comparisons (**Supplementary Figure S2D**).

**Figure 2.**
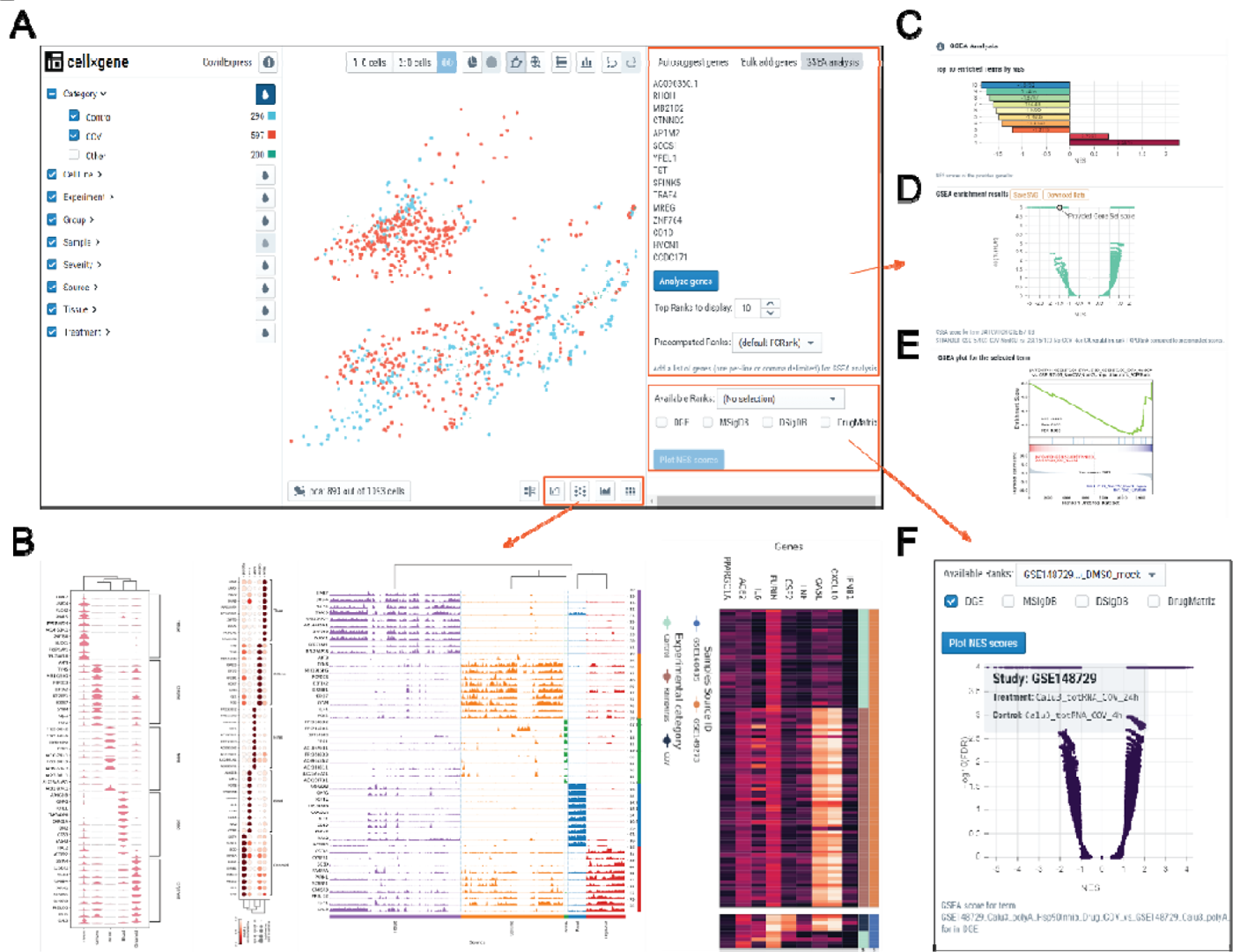
CovidExpress web portal overview and key functional components. **A** Main Graphical User Interface of CovidExpess web portal. Left panel were organized by meta data. Middle panel each dot represents a sample. Right panel allow user input gene of interest for reviewing their expression level or perform GSEA analysis. **B** Our newly designed visualization tool allow user generating violin plot, dot plot, track plot and heatmap for selected genes across all or selected samples. **C** Barplot of top GSEA results compared to pre-computed GSEA results. User mouse hover each bar would show detailed information. User click on bar would update Volcano plot and GSEA plot as panel D and E. **D** Volcano plot for the selected result. User mouse hover each dot would show detailed information. Data and plot could be downloaded for user’s customization. **E** GSEA plot for the selected result. User click could find more functions such as zoom, rotate, download. **F** Volcano plot for pre-computed data. Multiple choices of rank types and gene set databases available.

### Examples Demonstrating How to Use CovidExpress

CovidExpress is a rich resource for investigators at various stages of their research projects. For investigators who do not know exactly where to start the analysis in our web portal, we compiled a gene name cloud to highlight genes that have been frequently studied or implicated in SARS-CoV-2 and COVID-19 studies based on RNA-seq analysis (**Figure 1C, 1D, Supplementary Table S2**), and the functional significance of many of these transcripts remains to be determined. To showcase the utility of CovidExpress in exploratory analysis, we present below two case studies.

As the first case study, we wish to illustrate how CovidExpress portal might be used to investigate the expression patterns of pre-selected genes of interest and identify new ones. Here as the example, we examine the expression of a COVID-19 severity related gene set, identified by Overmyer et al (Overmyer *et al*, 2021), using an elastic net machine learning approach to find molecular features with high significance to COVID-19 status and severity based on multi-omics data. We restricted our analysis to 20 genes with the highest predictive power from the original paper. Then, using CovidExpress, we checked the expression of these genes in our compiled gene expression data of all the studies that reported the patient’s COVID-19 disease severity (510 samples from 14 different studies). The violin plot in **Figure 3A** indicates that, indeed, many of these genes show some relatively higher expression especially in the samples from Intensive Care Unit patients (ICU), Non-ICU, Remission and Severe patients. To closely check the enrichment pattern of these genes and avoid batch effect, we selected the ICU and Non-ICU samples from Overmyer et al. data (GSE157103) and plotted their expression in all of the 126 samples (66 ICU and 60 Non-ICU) (**Figure 3B, 3C**). The expression pattern of these genes indicates that *GRB10, ARF1, PGS1, RASGEF1A* and *SESN2* genes are highly expressed in ICU samples, while the rest of the genes are highly expressed in Non-ICU samples. These results can be very useful to get a sense about the role of these genes in COVID-19 severity.

**Figure 3.**
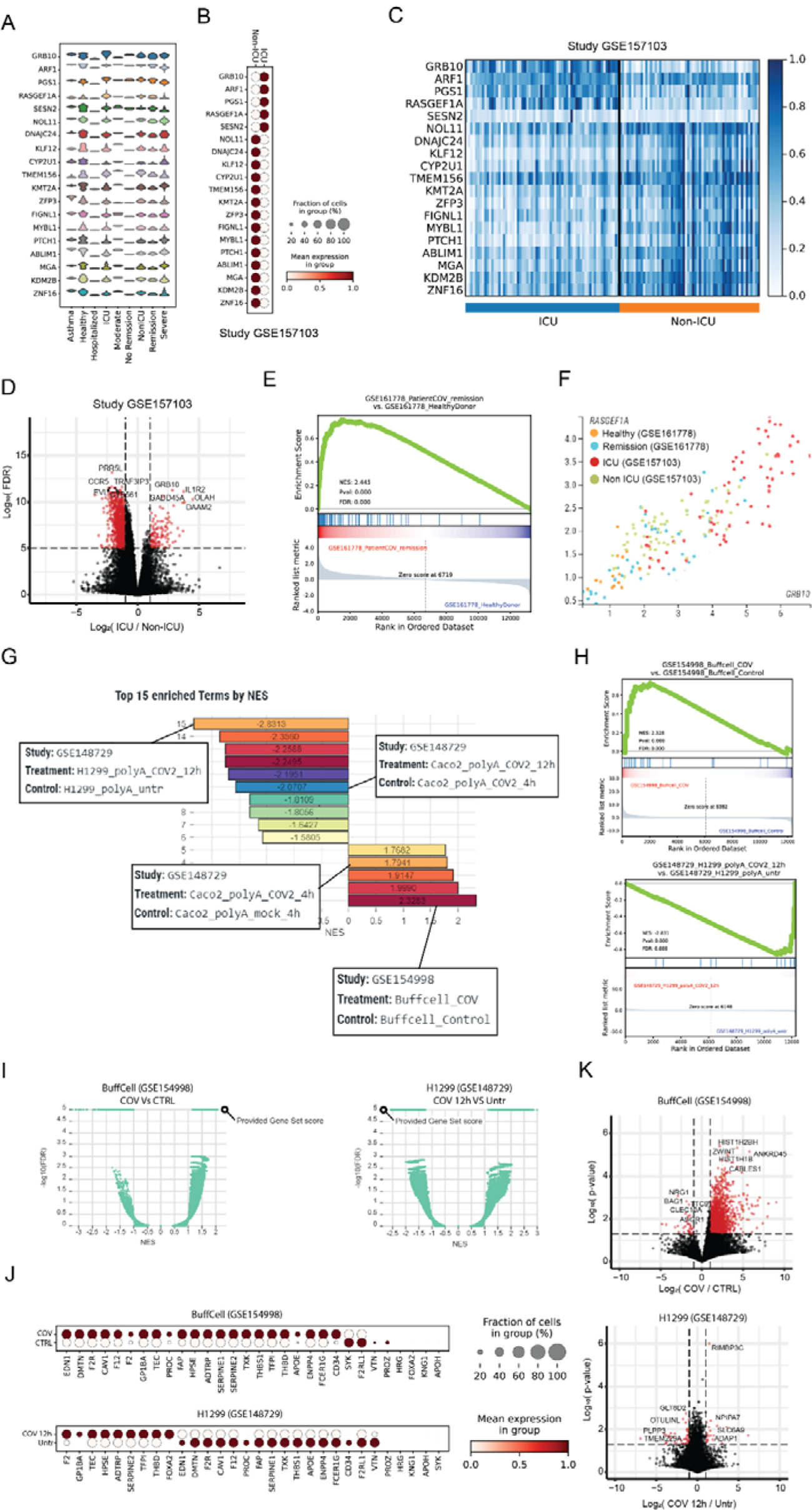
Use cases demonstrating the common steps for using CovidExpress. **A** Use case1: CovidExpress violin plot showing the expression of the top 20 COVID-19 severity predictors as defined by Overmyer et al (Overmyer *et al*., 2021) in the available severity classes available in CovidExpress. **B** CovidExpress dot plot showing the expression pattern of the top 20 COVID-19 severity predictors in Overmyer et al dataset (GSE157103). **C** CovidExpress heatmap showing the detailed expression of the top 20 COVID-19 severity predictors in Overmyer et al dataset (GSE157103). **D** Volcano plot showing the transcriptome-wide differential expression results as calculated by CovidExpress of ICU vs Non-ICU samples from Overmyer et al dataset (GSE157103). **E** CovidExpress GSEA plot showing the enrichment the upregulated genes from the ICU samples of GSE157103 in the remission patients from GSE161778. **F** CovidExpress scatter plot showing the gradual distribution of samples based on their COVID-19 severity using two top COVID-19 severity predictors. **G** Use case2: CovidExpress GSEA ranking plot of the list of experiment showing the most enriched and depleted expression of the coagulation genes (GO:0030193). **H** CovidExpress GSEA plot of the experiments showing the most enrichment and depletion of the expression of the coagulation genes. **I** CovidExpress GSEA volcano plot comparing the NES and FDR scores of the coagulation geneset relatively to the other contrasts in the samples identified in figure H. **J** CovidExpress dot plot showing the expression pattern of the coagulation genes in the experiments identified in panel H. **K** Volcalo plot showing the identification of additional differential genes between the previous two experiments using CovidExpress differential gene expression feature.

To investigate further the difference between ICU and Non-ICU patients, we performed differential gene expression analysis using CovidExpress for the same study (GSE157103). We identified the significantly up- and down-regulated genes between these two conditions and found some gene from the starting list also were among top differential expressed genes such as *GRB10* (**Figure 3D**). The results of GSEA analysis run for the ICU up-regulated genes indicates that the ICU enriched genes are a good discriminator between healthy controls and patients in the remission state from another study (GSE16778) further supporting the importance of these genes (**Figure 3E**). Consistently, the expression for top COVID-19 severity predictor genes *GRB10* and *RASGEF1A* were correlated and were overall higher in both ICU vs. non ICU and remission vs. healthy patients (**Figure 3F**).

As the second case study, we wish to illustrate how investigators can utilize CovidExpress to explore dozens of datasets starting from biological hypothesis and ending on an in-depth analysis of selected studies. An increasing body of literature is showing that altered coagulation is one of the strong phenotypic markers associated with severe COVID-19 cases (Al-Samkari *et al*, 2020; Osuchowski *et al*, 2021; Ramlall *et al*, 2020). Thus, we wanted to check how genes related with blood coagulation (GO:0030193) are regulated in different samples in our cohort. Using the GSEA enrichment feature, we can first identify the top 15 samples that show a significant up- or down-regulation of the coagulation markers (**Figure 3G**). Among the contrasts that show an up-regulation of the coagulation genes are the Buffy coat cells from COVID-19 positive ICU patients compared with control (GSE154998); while on the other hand, the SARS-CoV-2 treated H1299 cells (GSE148729) showed decreased expression of coagulation genes after treatment. Interestingly, in Wyler et al. study (Wyler *et al*, 2021) H1299 cell lines showed a low susceptibility to SARS-CoV-2 infection due to their low expression of ACE2 and low virus replication. This in turn might explain the altered activation of the coagulation genes.

To examine the degree of up/down-regulation of our coagulation gene set, we can further explore the GSEA enrichment profile. Indeed, most of the genes are up- and down-regulated in GSE154998 and GSE148729 experiments, respectively (**Figure 3H**). We can also compare how the GSEA score calculated for our coagulation gene set are ranked in respect with the precalculated contrasts from our database. For example, **Figure 3I** illustrates volcano plot, for the relation between Normalized Enrich Score (NES) and p-value, for the Buffy coat cells (left) and H1299 cells (right). Using the available custom annotation feature, investigators can plot the gene expression of the different coagulation genes (**Figure 3J**) and perform differential gene expression analysis to study in more details the expression patterns of these experiments (**Figure 3K**). As expected, and as reported by the previous studies, Buffy coat cells shows a large change in its genome-wide expression, while the H1299 cell line showed a transcriptional profile similar to uninfected cells as reported by Wyler et al. (Wyler *et al*., 2021). We summarized suggested analysis steps for various investigation interests in **Supplementary Figure S3A**. Overall, CovidExpress provided functionalities can become a very handy and powerful exploratory tool for investigators, especially those without advanced programming background or without easy access to high-performance computing facilities.

### CovidExpress Reveals Insights and Potential Discoveries

With all re-processed datasets and web portal in CovidExpress, we sought to explore user friendliness of the web portal. The first challenge that we encountered was the heterogeneity of experimental protocols and data variations. As expected, samples clustered strongly by study cohorts in conventional PCA (**Figure 4A**). PCA reduction using the top 1000 differentially expressed genes leads to some improvements, but the batch effect was not totally removed (**Supplementary Figure S4A).** Different dimension reduction methods such as t-distributed stochastic neighbor embedding (tSNE) or Uniform Manifold Approximation and Projection (UMAP) (McInnes *et al*) did not solve this problem, either (**Supplementary Figure S4B,C)**. Different scaling options such as batch correction using Combat (Leek *et al*, 2012) could not entirely solve this problem (**Supplementary Figure S4D).** We hypothesized that the inherent experimental variability might be partially mitigated or equalized by correcting against a set pathways or regulated genes using the approach of Single-sample GSEA (ssGSEA) enrichment scores (Barbie *et al*) for PCA, tSNE and UMAP analysis instead of correcting at the gene level. ssGSEA is an extension of GSEA to calculate separate enrichment scores for each pairing of a sample and gene set. Each ssGSEA enrichment score represents the degree to which the genes in a particular gene set are coordinately highly or lowly expressed within a sample. By calculating ssGSEA in each sample, we were able to get an enrichment score for each gene set in MSigDB (Liberzon *et al*.), including well known pathways like KEGG (Kanehisa *et al*), REACTOME (Jassal *et al*), BIOCARTA (Nishimura), Wikipathways (Slenter *et al*), and Gene Ontology processes (The Gene Ontology). Next, we used ssGSEA scores to cluster samples by similarity between gene sets, with the rationale that ssGSEA scores are sufficient to represent gene variability between samples.

**Figure 4.**
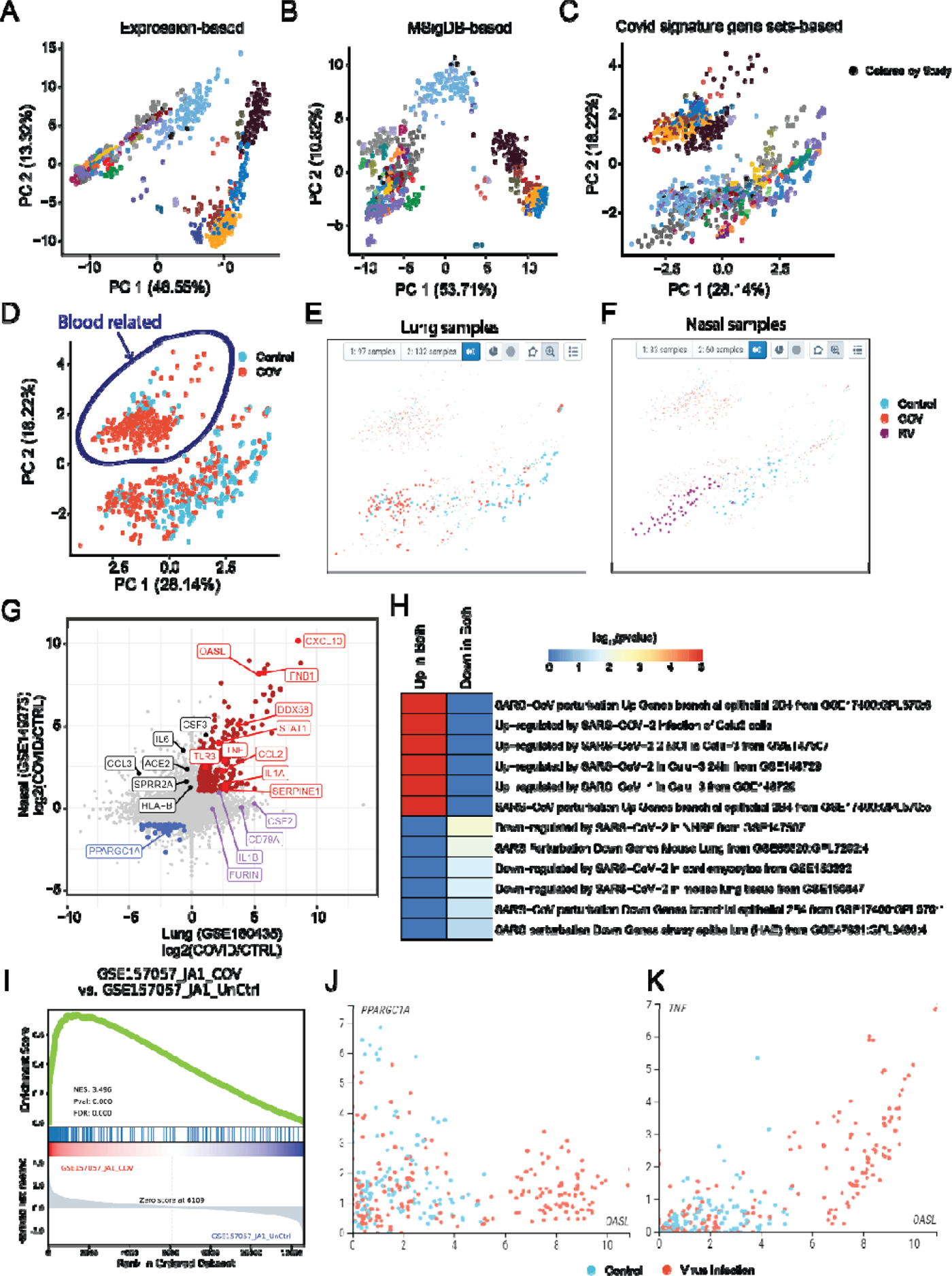
CovidExpress reveal insight led to potential discovery. **A** PCA analysis based on gene expression level (TPM), colored by studies **B** PCA analysis based on single-sample Gene Set Enrichment Analysis(ssGSEA), using gene sets from MSigDB, colored by studies **C** PCA analysis based on single-sample Gene Set Enrichment Analysis(ssGSEA), using Covid signature gene sets from this study, colored by studies **D** PCA analysis based on single-sample Gene Set Enrichment Analysis(ssGSEA), using Covid signature gene sets from this study, colored by SARS-CoV-2 infection status. **E** Using combination of lasso selection and meta data selection, we selected lung tissue SARS-CoV-2 (red dots) and control group (blue dots). **F** Using combination of lasso selection and meta data selection, we selected nasal tissue Rhinovirus (red dots) and control group (blue dots). **G** Scatterplot showing the log_2_ expression fold change between lung and nasal cells after virus infection. Genes up-regulated in both tissues are shown in red, while the down-regulated in both tissues are shown in blue. We labelled genes shown as top in our literature-based word cloud that black ones were only significant up-regulated by Rhinovirus in nasal, while purple were only significant up-regulated by SARS-CoV-2 in lung. **H** Heatmap showing the -log_10_(p-value) of the commonly up-/down-regulated genes in COVID-19 genes sets compiled by the EnrichR database. **I** GSEA plot of one of top contrast differential expression comparison (GSE157057) showing the enrichment of the commonly up-regulated genes in lung and nasal tissue. **J** Scatterplot showing the anti-correlation between *OASL* (commonly up-regulated) and *PPARGC1A* (commonly down-regulated) genes in nasal and lung samples. Samples are colored based on their phenotype (red: SARS-CoV-2 or Rhinovirus, blue: Control). **K** Scatterplot showing the correlation between *OASL* and *TNF* (both commonly up-regulated) in nasal and lung samples. Samples are colored based on their phenotype (red: SARS-CoV-2 or Rhinovirus, blue: Control).

To test this hypothesis, we used GTEx data as a ground truth. We downloaded and processed 9,525 GTEx samples from 30 tissues, then, we calculated the top 10 principal components projection for each sample using its gene expression and MSigDB ssGSEA enrichment scores respectively (**Supplementary Figure S4E, S4F**). Next, we used the silhouette score to measure the separability between tissues (Rousseeuw, 1987) and found that ssGSEA scores-based projection indeed leads to a better separability between tissues (**Supplementary Figure S4G**).

Encouraged by these results, we then applied the ssGSEA approach on our data collection, we observed that the samples clustered less according to study cohorts (**Figure 4B, Supplementary Figure S4H**). This clustering was further improved if we use the ssGSEA score from COVID-19 signature gene sets (top differentially expressed genes from our analysis) (**Figure 4C, Supplementary Figure S4H**). In contrast, although batch effect correction method such as Combat achieved the best experiments-based silhouette score (i.e. making samples less separated by study cohorts)(**Supplementary Figure S4H**), this strategy did not improve the samples separation by phenotype (infected with SARS-CoV-2 or not)(**Supplementary Figure S4I**). As expected, this clustering did not perfectly separate infected samples from control samples (**Figure 4D**) due to tissue specificity and many infected samples were also tested with drug treatments. In the portal, we show a projection obtained by using the ssGSEA scores of the differentially expressed gene sets by default setting in the CovidExpress portal, but investigators may select other types of projections (**Supplementary Figure S5A, S5B, S5C**).

Next, we utilized the portal’s function to gain insights about SARS-CoV-2’s effect. We first noticed there were two large clusters despite their SARS-CoV-2 infection status (**Figure 4D**). Interestingly, the left top cluster were all from blood related samples such as white blood cells, leukocytes, monocytes. While the right bottom cluster were from other tissues. This indicated the COVID-19 signature was not able to overcome the tissue specificity and that immune cells might respond to SARS-CoV-2 differently. Next, we reviewed the SARS-CoV-2 infection status for studies with large number of samples. CovidExpress portal allowed us to select different projection of samples (**Supplementary Figure S5A**), that we noticed Rhinovirus samples separated from control samples but mixed with some SARS-CoV-2 samples in both UMAP and tSNE projections (**Supplementary Figure S5B, S5C**). The clustering of Rhinovirus samples is less visually impressive in PCA. However, we noticed PC1 generally reflected the effect of virus infection and that smaller PC1 value correlated with virus infection. For example, more SARS-CoV-2 infection samples clustered at left in lung (**Figure 4E**), similarly more Rhinovirus (RV) infection samples clustered at left in nasal sample (**Figure 4F**). This is consistent with the proposed role that RV infections are potential mechanisms of ACE2 overexpression in patients with asthma (Chang *et al*, 2020). We next selected one study from each tissue that had a simple experimental design comparing SARS-CoV-2 or RV infected cells versus control (GSE160435 and GSE149273) and identified their differentially expressed genes respectively.

Interestingly, 345 genes (280 up-regulated and 65 down-regulated) showed consistent expression change pattern in both lung and nasal cells (**Figure 4G, Supplementary Table S3**). Among the 280 consistently up-regulated genes, 56 genes (20%) where among the top 1000 genes highly cited in COVID-related literature mined through the LitCovid database (Chen *et al*., 2020, 2021a) (**Figure 4G**). Among the top hits we identified genes such as *TNF*, *IL1A* and *CXCL10* previously identified as a common factor between pulmonary and olfactory dysfunctions in SARS-CoV-2 infections (Oliviero *et al*, 2020). Additionally, TNF has been functionally shown, along with IFN-γ, to be one of the key drivers of cytokine storm in COVID-19 (Karki *et al*, 2020). Functional enrichment analysis against the COVID-19 gene-sets compiled by the EnrichR database (**Figure 4H**) (Kuleshov *et al*., 2020) shows up/down-regulation patterns consistent with those from other independent studies. These enrichment results also indicate that the identified up/down-regulated genes are also consistently changing in the airway epithelium cells (**Figure 4H**). GSEA analysis of the identified up-regulated genes against the gene contrasts compiled in our database also indicate consistent upregulation in ex vivo cells such as in SARS-CoV-2 infected lung organoids (**Figure 4I**).

Among the commonly differential genes in both lung and nasal tissues we also identified an interferon response gene *OASL*, an under-studied protein from oligoadenylate synthase (OAS) protein family. OASL’s role in antiviral activity has been reported that it could enhance RIG-I (Zhu *et al*, 2014) but it was not extensively studied in the context of COVID-19 based on LitCovid database. Interestingly, we observed that, *OASL* showed an inversely correlated pattern to the common down-regulated gene such as *PPARGC1A* and can overall discriminate the lung and nasal virus infected cells from the control group (**Figure 4J**). The expression of *OASL* is also highly correlated with that of *TNF* in COVID samples (**Figure 4K**). Among all tissues, *OASL* expression is highest in the lung tissue (**Supplementary Figure S5D**), particularly in macrophage cells in lung tissue (**Supplementary Figure S5E**), suggesting its important role in the innate immune response in lungs.

We also noticed *ACE2* and several other top studied genes were significantly up-regulated only in RV infected nasal but not in SARS-CoV-2 infected lung organoids (**Figure 4G**). This might also be due to the nature of SARS-CoV-2 needing ACE2 to infect cells (Clausen *et al*.; Hoffmann *et al*., 2020; Liu *et al*., 2020; Ziegler *et al*., 2020) but not regulating *ACE2*’s expression. Interestingly, despite “IL-6/JAK/STAT3 Signaling” were enriched for genes up-regulated in both lung and nasal (**Supplementary Table S3**), *IL6* is significantly up-regulated only in RV infected nasal but not in SARS-CoV-2 infected lung organoids (**Figure 4G**). This indicated the activation of IL-6 signaling were different between RV infected nasal and SARS-CoV-2 infected lung organoids. In lung, there are other pathways implicated such as mTOR (**Supplementary Table S3**) (Mullen *et al*, 2021). In addition, furin cleavage site in SARS-CoV-2 was reported to help its infectivity and transmissibility (Johnson *et al*, 2021; Xia *et al*, 2020) while our results found *FURIN* gene was also up-regulated in SARS-CoV-2 infected lung organoid alveolar type 2 cell (**Figure 4G, Supplementary Table S3**) (Mulay *et al*, 2021), implicating a positive feedback, presumably via TGF-beta Signaling (**Supplementary Table S3**) (Blanchette *et al*, 2001). Given that the furin cleavage site could naturally occur in coronavirus family (Wu & Zhao, 2021), it is plausible SARS-CoV-2 might evolved to induce *FURIN* expression to gain superior infectivity, although this hypothesis requires further investigation. Together, the data analysis capacities offered by CovidExpress enable scientists to identify key genes and pathways that would be catalysts of new scientific investigations.

## DISCUSSION

We describe the data collection, data processing, and web portal development of a comprehensive RNA-seq database from SARS-CoV-2 and COVID-19 related research, named CovidExpress. CovidExpress database content and portal will be periodically updated. We expect the updates will be based on the framework described here. We also plan to improve our differential gene expression analysis component in the future to adjust for batch effects between studies. **For the current release, we strongly suggest investigators to perform gene expression comparison within individual study**. To make the database as user friendly as possible, we have developed abundant visualization functions to help investigators to quickly test their hypothesis and visualize results. We included GSEA component to enable investigators to analyze gene sets on-the-fly and compare results with thousands of pre-computed GSEA results. This framework could be easily applied to different collections of data. Finally, our approach in using ssGSEA scores for unbiased clustering sheds light on how to visualize large datasets with varying, strong, and most importantly unknown batch effect.

We strongly believe the CovidExpress database has a power to become a key tool for identifying new genes of interest to support the development of hypotheses and subsequent biochemical analyses. While many genes have been found to be differentially regulated during SARS-CoV-2 infection across studies, some are only found in specific cell types or experimental conditions, and the functional relevance of many of these remains unclear. It will be beneficial to use the CovidExpress tool as the first step in elucidating these functional pathways across dozens of experiments. Follow up investigations aimed at elucidating where genes of interest fit into biochemical pathways and characterizing which are upstream regulators or sensors that can affect a variety of downstream processes will be essential to improve our understanding of SARS-CoV-2 pathogenicity and identify new therapeutic approaches. For example, while IL-6 has been identified in many datasets as being important in COVID-19, biochemical characterization has lagged, and the functional relevance of IL-6 remains unclear, as clinical trials to block IL-6 or its receptor have had mixed results in patients (Gordon *et al*, 2021; Nasonov & Samsonov, 2020). Additionally, analyses should consider the stage of disease, which can be selected using the sample selection feature in CovidExpress. Innate immune responses, particularly IFN responses, are known to change throughout the infection (Blanco-Melo *et al*, 2020; Hadjadj *et al*, 2020; Lee *et al*, 2020; Lee & Shin, 2020; Lucas *et al*, 2020), making temporal considerations key.

Overall, the CovidExpress database fills a critical gap in the research field to comprehensively compile the large amount of RNA-seq data available on COVID-19 and offer it in a format that enables users to perform basic and visualizable analyses without the need for programming skills. We think this work will not only benefit SARS-CoV-2 research field but may also inspire other data-driven investigators on how to utilize rich data already published for scientific discoveries.

## DATA AVAILABILITY

RNA-seq data were collected from GEO as listed in **Supplementary Table S1**. Raw data with annotation and gene signatures could be downloaded from https://stjudecab.github.io/covidexpress

## MATERIALS AND METHODS

### RNA-seq data analysis

Sequencing reads were quality filtered using TrimGalore (available on-line at https://www.bioinformatics.babraham.ac.uk/projects/trim_galore/). Filtered reads were aligned to the human reference genome GRCh38.p12 using STAR (Dobin *et al*, 2013), assuming that the RNA-seq experiment is strand-specific. Next, MarkDuplicates from GATK (McKenna *et al*, 2010) and CollectRnaSeqMetrics from Picard (available on-line at http://broadinstitute.github.io/picard/) were used to mark duplicated reads and compute mapping statistics. Only samples with more than 2M reads mapped (1,460 samples) or more than 1M deduplicated reads mapped (1,396 samples), were later retained for further analysis.

RSEM (Li & Dewey, 2011) was used to quantify read counts per gene based on Gencode v31 reference gene annotation, and expression values were converted to Transcripts Per Kilobase Million (TPM) unit. Next, infer_experiment.py tool from RSeQC (v4.0.0) (Wang *et al*., 2012) was used to examine strand-specificity for each sample. Subsequently, the results from each sample were manually examined to determine final strand-specificity. If majority of samples in experiment (i.e. samples from the same GEO accession number) exhibited no bias toward strand specificity, as emphasized by fraction of reads mapping to both strands at the level > 0.4 and < 0.6, the experiment strategy was changed to unstranded (non-strand specific), and reads from all samples from that experiment were remapped in the unstranded mode. The exception from the approach to assign the same strand specificity toward all samples from the same experiment was made for GSE147507 experiment, in which case the manual examination revealed that strand specificity status called by RSeQC was consistent within subseries of the samples, as emphasized by the sample names. E.g. samples from “Series1” were consistently considered strand-specific, while samples from “Series2” were non-strand specific.

After all samples were mapped, geneBody_coverage2.py from RSeQC was used to examine the percentile-distribution of mapping reads along housekeeping genes. Distribution in each sample was min-max normalized, hieratically clustered with Ward’s method and manually examined in order to subjectively identify clusters representing samples with the highest quality (clusters #4, #7, #8, and #9; 946 samples), medium quality (clusters #5 and #6; 163 samples) and low quality (clusters #1, #2 and #3; 359 samples; **Supplementary Figure 1H**). Normalized distributions were also used to calculate for each sample the cumulative sum at 50^th^ percentile, with the assumption that deviation of this cumulative value from 0.5 value will increase with the increasing bias toward mapping of reads at 3’ or 5’ end. The lower and upper thresholds to consider a sample as being biased toward 3’ or 5’ end was set as 0.2716 and 0.7496 respectively; which values corresponds with the minimal and maximal values of cumulative sum at 50^th^ percentile (min=0.3282; max=0.693), minus/plus one standard deviation (std=0.0566), computed among 946 samples which were considered as representing the highest quality gene coverage among studied samples. Applying those thresholds for all samples allowed for identification of total of 1,274 samples with low 5-3 ends bias. Independently, the mean coverage was calculated for each sample from normalized gene body coverage distributions, with the assumption that the value will be lower for low quality samples than for high quality samples. Indeed, for example the mean coverage for 359 samples from three lowest quality clusters #1, #2 and #3 (**Supplementary Figure 1H**) the mean coverage is on average at the level of 0.2669, while in contrast for 946 samples from high quality clusters #4, #7, #8, and #9, this value is at the level of 0.7392. Therefore, a value 0.3843 was set as a minimal average gene body coverage threshold, which value corresponds with the minimal coverage value of 0.5138 (minimal among 946 highest quality samples) minus three standard deviations from the mean (std=0.04317). This allowed for identification of 1,160 samples meeting average gene body coverage threshold. Finally, carefully considering all quality control criteria, a total of 1,093 samples were selected for further analysis (**Supplementary Figure 1J**).

After removing low quality samples, batch correction using Combat (Leek *et al*., 2012) was applied for FPKM values, which values were used only for the purpose of samples clustering with PCA, if the user would like to choose this option. However, because of the nature of the CovidExpress project, and very large technical and biological variability between experiments, in order to minimize the influence of the batch effects correction, the differential gene expression analysis was computed only for the samples originating from the same experiment based on non-batch-corrected FPKM expression values. Differential expression analysis was assessed using limma-voom (Law *et al*, 2014). Only GENCODE annotated level 1 and 2 protein-coding genes, with at least 10 reads per sample in the minimum group size, were retained in the analysis. Based on each contrast, differential genes were extracted with the following thresholds:

1. Up2: FC ≥ 2, FDR ≤ 0.05
2. Up2NoFDR: FC ≥ 2, p ≤ 0.05
3. Up: FDR ≤ 0.05
4. UpNoFDR: p ≤ 0.05
5. DownNoFDR: p ≤ 0.05
6. Down: FDR ≤ 0.05
7. Down2NoFDR: FC ≥ 2, p ≤ 0.05
8. Down2: FC ≥ 2, FDR ≤ 0.05

Moreover, based on each DEG contrast, the genes were pre-ranked following four alternative approaches:

1. FCRank metric = log_2_ (FC)
2. PRank metric = direction × -log_10_ (p-value)
3. FCPRank metric = log_2_ (FC) × -log_10_ (p-value)
4. FCPERank metric = log_2_ (FC) × -log_10_ (p-value) × log_10_ (Mean Expr. +1)

Each pre-ranked gene list was sorted and used to extract 20, 50, 100, 200, 500, 1000 and 2000 Top (upregulated) and Bottom (downregulated) genes. All pre-ranked gene lists were also used to calculate Gene Set Enrichment Analysis (Subramanian *et al*., 2005) using GSEApy (v.0.10.2, available online at https://github.com/zqfang/GSEApy). GSEApy was run with 1000 permutations and gene set size thresholds were set to 5 and 5000 for minimal and maximum size, respectively. The analysis was independently conducted for gene signatures collections from four sources:

1. COVID-19 related gene signatures: 561 differential gene signatures from COVID-19 related research, downloaded from Enrichr portal (Kuleshov *et al*, 2016; Kuleshov *et al*., 2020). This collection was enlarged by 20,036 differential gene signatures collected at various thresholds and top up-/down-regulated genes from CovidExpress project.
2. DSigDB: 23,950 gene signatures from DSigDB database (Yoo *et al*, 2015), downloaded through Enrichr database.
3. DrugMatrix: 7876 gene signatures from DrugMatrix database (Ganter *et al*, 2006), downloaded through Enrichr database.
4. MSigDB: 25,724 gene signatures from MSigDB (v7.1) (Liberzon *et al*., 2015; Liberzon *et al*, 2011). In addition to pre-ranked GSEA analysis, single-sample GSEA, also using GSEApy program, was conducted. Normalized enrichment scores (NES) were computed for all the COVID-19 related samples either for the full collection of gene signatures from MSigDB, or collection of in-house differential gene signatures from COVID-19 related research, narrowing down the full collection to 492 differential gene signatures from Up2 and Down2 categories.

### CovidExpress web portal development

We built CoivdExpress by leveraging the sophisticated visualization features of cellxgene interface, which is a tool that was originally designed for exploring single-cell RNA-seq analysis. Because cellxgene client side was built using the React library, the addition of new user interface (UI) components can be done in a straightforward modular way. We mainly added the following components: GSEA analysis and visualization related components, gene expression visualization plots and differential gene expression analysis and full results downloading component. We used nivo library (https://nivo.rocks/) for visualization of the interactive plots. The GSEA enrichment plot was generated using the python module gseapy (https://github.com/zqfang/GSEApy), while the different gene expression plots were generated using the python module scanpy (Wolf *et al*, 2018).

### Data preparation for visualization

To be able to pass the data to cellxgene, it needs to be converted into the .h5ad format understood by scanpy (Wolf *et al*., 2018). Thus, we first used Seurat v3 R package (Stuart *et al*, 2019) to load the expression data, add the different projections slots and the metadata. The Seurat object was then converted into the .h5ad format using the SeuratDisk R package (https://github.com/mojaveazure/seurat-disk).

The GSEA ranks, NES scores and p values, on the other hand, were stored in a separated .h5 file for rapid access and parsing.

## Supporting information

Table.S1.QC_metadata_contrasts

Table.S2.WordCloud

Table.S3.DE_MSigDB_Combined

## ACKNOWLEDGEMENTS

We thank High Performance Computing Facility at St. Jude Children’s Research Hospital for technical support. We thank M. Madan Babu, PhD, FRSC for his kindly feedback on improving the readability.

---

**Figure S1.**
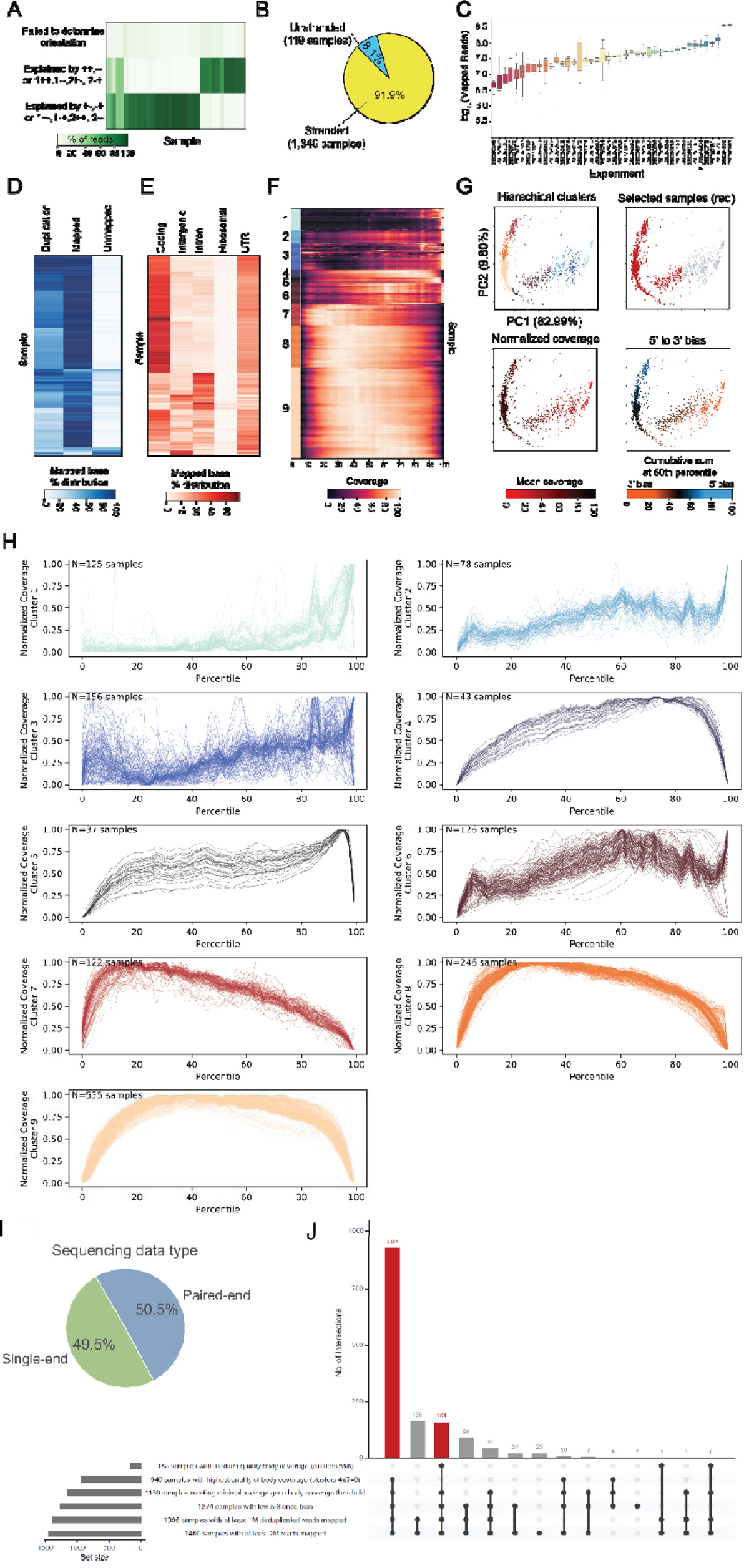
**A** Heatmap of results from automatic detection of RNA-seq strandness. **B** Pie chart shown percentage of stranded RNA-seq data versus unstranded. **C** Boxplot of mapped reads number by studies. **D** Heatmap of mapping rate, duplication rate. **E** Heatmap of read genomic distribution. **F** Hierarchical clustering of read aggregative gene-based coverage. **G** PCA plot colored based on hierarchical clusters (in panel G), 5’ to 3’ bias score and Average normalized coverage. Number in brackets indicated variance explained. **H** Line plot of aggregative gene-based coverage for individual cluster (defined in Supplementary Figure 1F). **I** Pie chart shown the percentage of RNA-seq sample using single-end versus paired-end sequencing. **J** Upset plot summaries the overlap between different clusters, groups and quality control criterions. The black dot in bottom right matrix indicated which group the sample for that bar belong to. For example, the 166 samples for second bar were overlapping only between group 1396_samples_with_at_least_1M_deduplicated_reads_mapped and 1460_samples_with_at_least_2M_reads_mapped.

**Figure S2.**
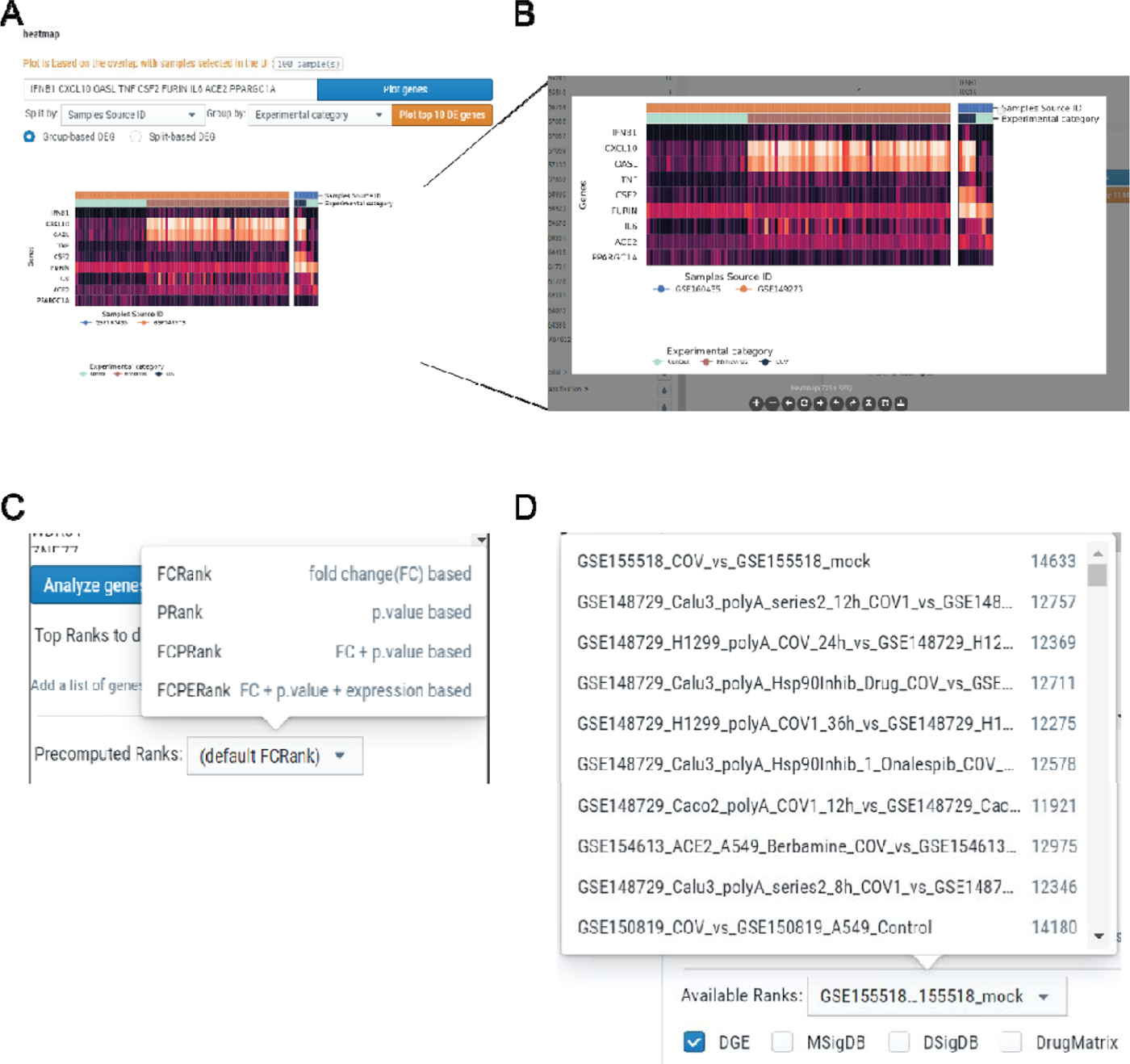
**A** Detailed demonstration of plot function. These functions were made available for all types of plots as violin plot, dot plot, track plot and heatmap plot. User could input their gene list. User could also simple plot the top 10 differential expressed gene by selected meta data (“Source” as example) here. Results here indicated very few genes could be distinct one source from the others. For selected group 1 versus group2, differentially expressed genes between group 1 and group 2 could also been plot. **B** Plot customization functions were provided so user could zoom, rotate, change text size, download the figure on-the-fly. **C** We provide GSEA for different options of pre-computed ranks. User could choose based on different questions to ask. For example, using fold change (FC) based rank if they wondering what have been enriched by most changed genes. p.value based if they more interesting in what have been enriched by most consistently changed genes. See Material and Methods part for details. **D** An example of how pre-computed results were organized and available for user’s selection.

**Figure S3.**
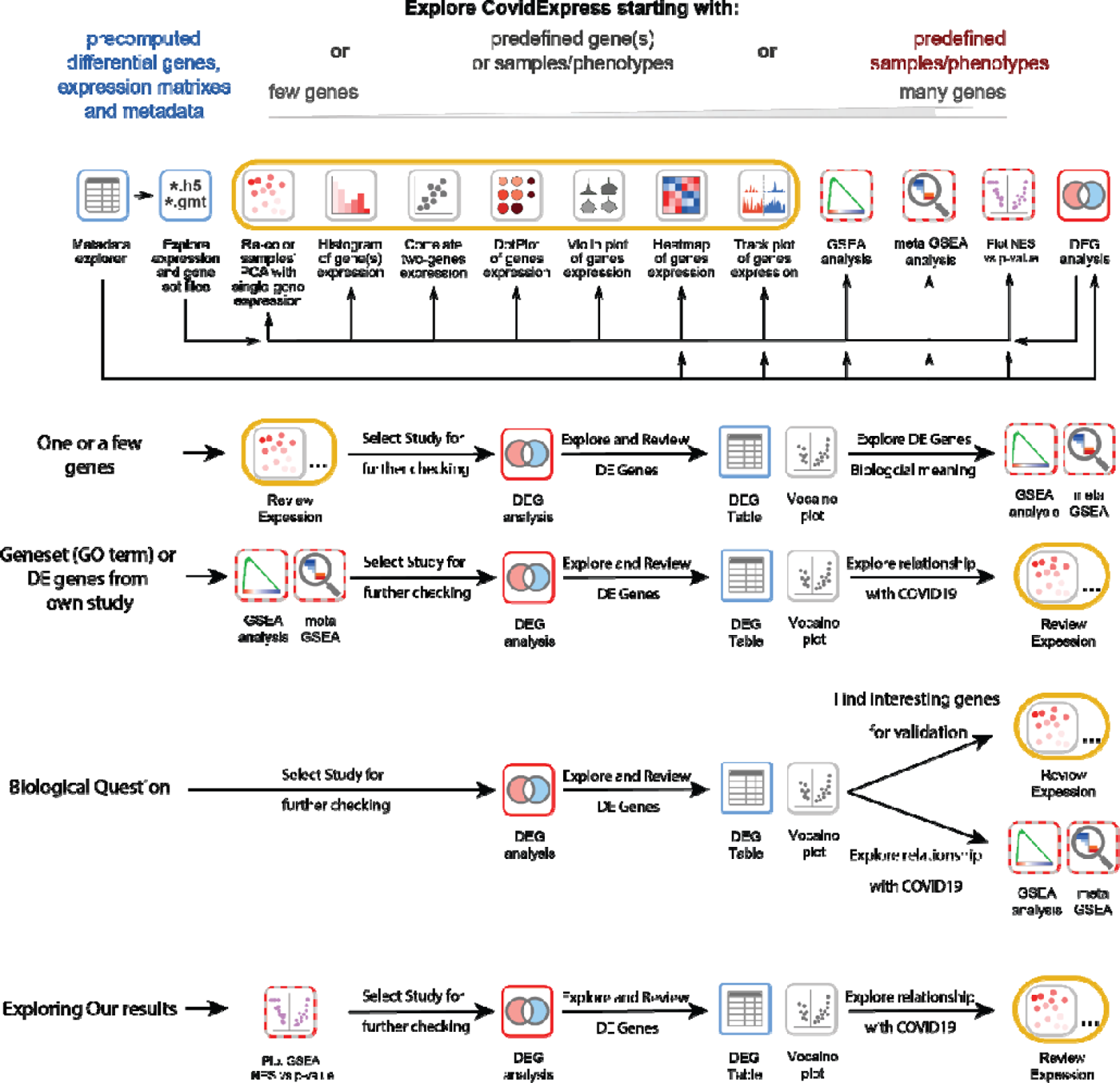
Suggested analysis steps for variously investigation interests.

**Figure S4.**
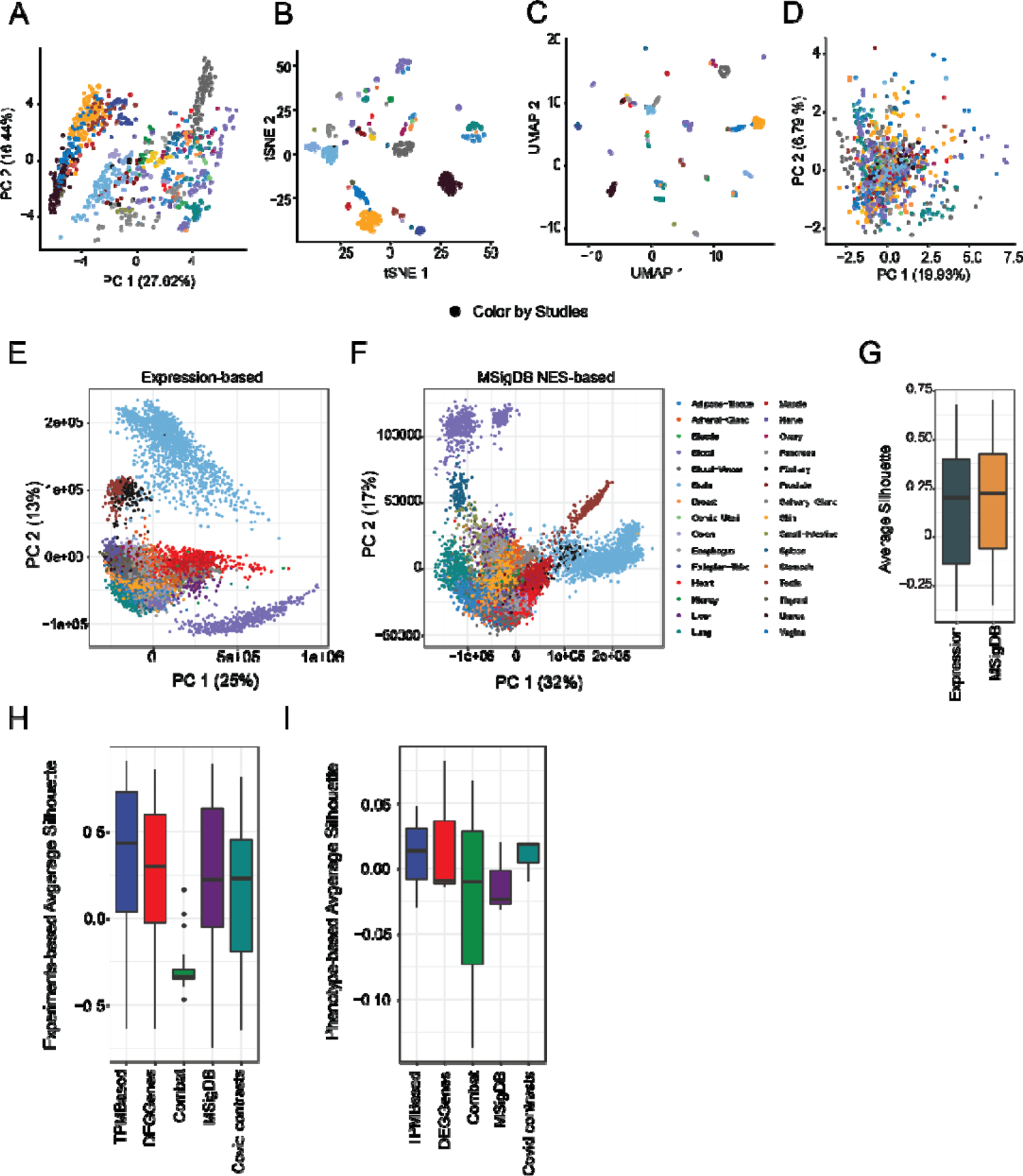
**A** PCA analysis based on the top one thousand differential genes, colored by studies **B** tSNE analysis based on gene expression level, colored by studies **C** UMAP analysis based on gene expression level, colored by studies **D** PCA analysis based on gene expression level after batch correction, colored by studies **E** PCA analysis of GTEx data based on gene expression, colored by tissue. **F** PCA analysis of GTEx data based on MsigDB signature, colored by tissue. **G** Comparison of tissue-level Silhouette score distribution using expression and MSigDB-based PCA projections. **H** Experiment-level Silhouette scores distribution between our compiled samples using different scoring methods to measure batch-effect. Lower score indicates less batch effect. **I** Phenotype-level Silhouette scores distribution between our compiled samples using different scoring methods to measure the degree of phenotype separability. Higher scores indicate better separability between SARS-CoV-2 infection and control samples.

**Figure S5.**
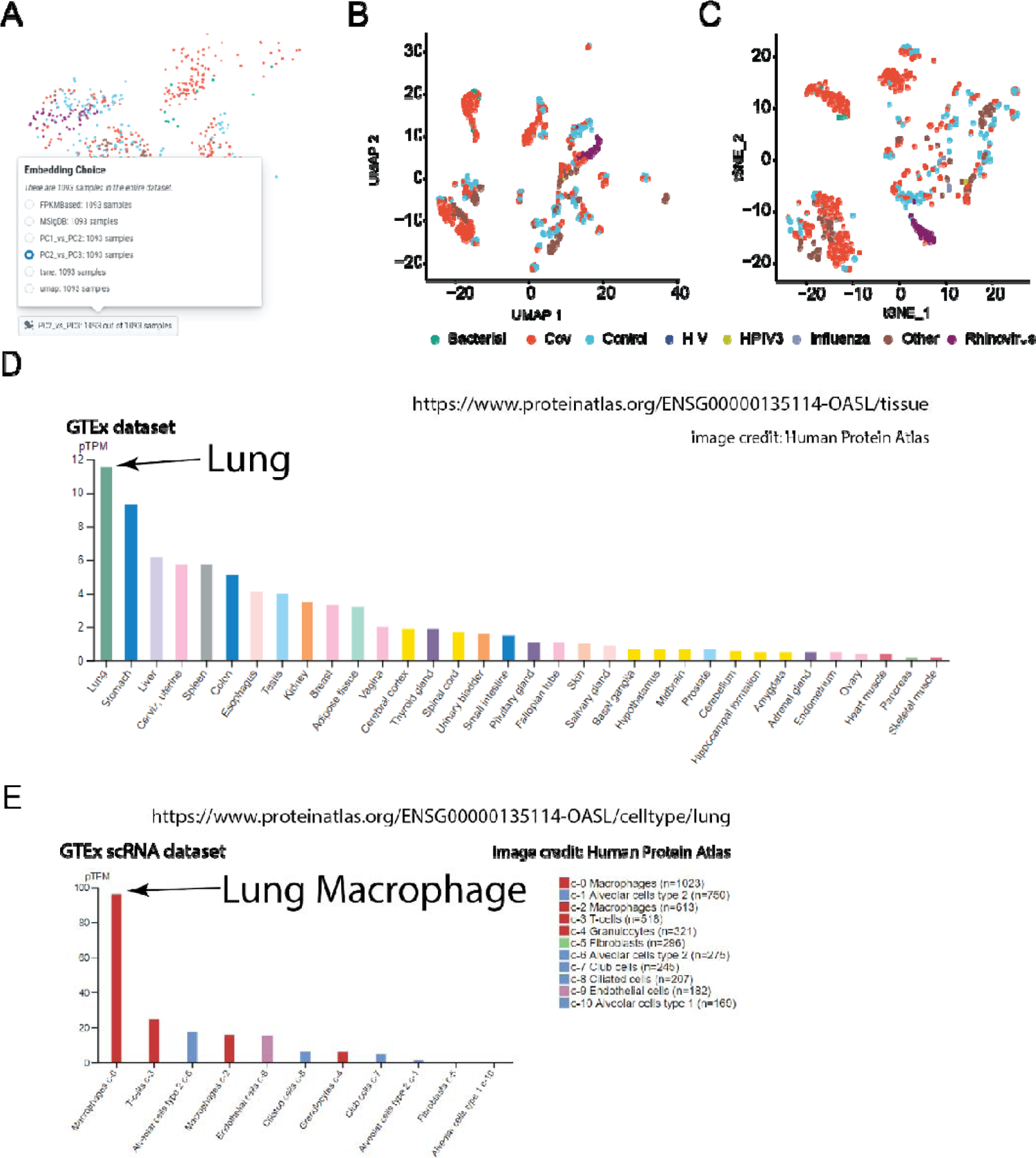
**A** An example shown various choices of samples embedding layouts user could use. **B** UMAP analysis based on single-sample Gene Set Enrichment Analysis(ssGSEA), using COVID signature gene sets from this study, colored by SARS-CoV-2 infection status. **C** tSNE analysis based on single-sample Gene Set Enrichment Analysis(ssGSEA), using COVID signature gene sets from this study, colored by SARS-CoV-2 infection status. **D** *OASL* expression in different tissues from GTEx datasets. **E** *OASL* expression in different cell types from Lung GTEx single-cell RNA-seq datasets.

